# Harnessing light-activated gallium porphyrins to combat intracellular *Staphylococcus aureus* in dermatitis: Insights from a simplified model

**DOI:** 10.1101/2023.12.13.571407

**Authors:** Klaudia Szymczak, Michał Rychłowski, Lei Zhang, Joanna Nakonieczna

## Abstract

*Staphylococcus aureus* can survive inside nonprofessional phagocytes such as keratinocytes, demonstrating a novel strategy for evading antibiotic pressure. When antibiotic treatment ends, reinfection with staphylococci begins from the intracellular inoculum. This phenomenon is responsible for recurrent infections. The development of new antibacterial methods that can eliminate intracellular bacteria, including those with a multidrug-resistant phenotype, is necessary. In this study, we characterized and used a model of keratinocytes (both wild type and mutants with reduced filaggrin expression) infected with methicillin-resistant *S. aureus* (MRSA) to verify the possibility of using light-activated compounds, exemplified here by heme-mimetic gallium (III) porphyrin (Ga^3+^CHP) and visible light, an approach known as antimicrobial photodynamic inactivation (aPDI), to eliminate intracellular MRSA. We observed that Ga^3+^CHP accumulated more in infected cells than in uninfected cells. Moreover, Ga^3+^CHP accumulated in cells that harbored intracellular *S. aureus*. Using flow cytometry and fluorescence microscopy, we found that intracellular MRSA and Ga^3+^CHP mainly colocalized in lysosomal structures, and we showed that under the influence of aPDI, MRSA exhibited reduced adhesion to host cells and a significantly reduced (by 70%) GFP signal originating from intracellular bacteria. Moreover, the use of light-activated Ga^3+^CHP resulted in a significant reduction in the number of extracellular bacteria in the infection system, lowering the potential for further infection of host cells. For the first time, we used the infectious model to analyze the toxicity of aPDI in real time, showing that this approach is not significantly cyto-or phototoxic.

**Author Summary:** *Staphylococcus aureus* is a highly virulent pathogen that is responsible for approximately 80% of all skin infections. During antibiotic treatment, one of the defense mechanisms of *S. aureus* is the invasion of keratinocytes. Intracellular bacteria are not accessible to antibiotics, which poorly penetrate the interior of host cells. Consequently, such bacteria contribute to recurrent infections. In our study, we proposed using a combination of a light-activated porphyrin compound loaded with gallium ions, Ga^3+^CHP, and visible light as a strategy to eliminate intracellular staphylococci. We demonstrated that the tested compound colocalized with the pathogen in the infected cells, which was an essential condition for the effective elimination of intracellular bacteria. We showed that the proposed approach effectively reduced the infection of keratinocytes with methicillin-resistant *S. aureus* (MRSA), as well as its adhesion to host cells, while maintaining host cells. The results presented here provide a basis for developing an effective therapy against staphylococci.

## Introduction

*Staphylococcus aureus* colonizes the skin in approximately 20% of the world’s population and is responsible for 80% of all detected skin infections worldwide (1,2). Moreover, *S. aureus* plays a key role in the pathogenesis of atopic dermatitis (AD) because its overexpression in the skin microbiota increases inflammation (3). Staphylococcal infections are difficult to treat due to the production of multiple virulence factors and a high antibiotic resistance profile (4). Despite antibiotic treatment, 30% of patients are reported to develop recurrent staphylococcal infections after the initial round of antibiotics (5). Recent scientific reports have shown that one of the defense mechanisms of *S. aureus* against antibiotic action is the invasion of nonprofessional phagocytes such as keratinocytes or fibroblasts (6–8). *S. aureus* internalization occurs through a complex zipper-like mechanism between fibronectin-binding proteins A and B and fibronectin, which is recognized by α5β1 integrin on host cells (9). The internalization process differs depending on the bacterial strain and host cell type (6). Under antibiotic pressure, *S. aureus* can persist inside host cells for several days as an intracellular form of bacteria known as small colony variant (SCV) bacteria (10). This phenotype exhibits changes in the global regulatory networks that can lead to alterations in the production of virulence factors or modify the response to antibiotics (11,12). When antibiotic pressure is abolished, the intracellular bacterium can escape the endosome and multiply in the cytoplasm. The increased number of bacteria inside the cell leads to cell death, release of bacteria, and recurrence of extracellular bacterial infection (13). Despite the increasing number of studies on *S. aureus*, the mechanism and factors contributing to the entry, survival, and exit of *S. aureus* from the host interior are still not fully understood.

The intracellular phenotype of *S. aureus* is caused and maintained by antibiotic action, as antimicrobials cannot effectively penetrate the cell membrane to achieve efficient intracellular killing (14). There are many proposed new therapies against intracellular *S. aureus* (15). Anti-intracellular bacteria strategies are based on antibiotic modifications to improve their delivery or cell stimulation that enhances bacterial killing by the host (16–19). Our study investigated antimicrobial photodynamic inactivation (aPDI) as a potential therapy against intracellular *S. aureus*. Its mechanism is based on three components: an oxygen-rich environment, light at the appropriate wavelength, and a small molecular weight compound with photodynamic properties – a photosensitizer (PS) (20,21). An ideal photosensitizer should exhibit a high reactive oxygen species (ROS) quantum yield with high phototoxicity against pathogens and low toxicity against eukaryotic cells. The penetration of PS into microbial cells should be rapid, with prolonged uptake in host cells (22,23). Gallium metalloporphyrins (Ga^3+^MPs) are potent PSs in aPDI that can absorb visible light, such as the green light used in this work (24–26). Ga^3+^MPs, due to their structural similarity to heme, can be recognized by bacterial heme-acquisition receptors of the Isd family and efficiently accumulate inside the bacterial cell (27,28). The porphyrin ring of Ga^3+^MPs is then cleaved inside the bacteria and gallium ions are released to disrupt iron-dependent metabolism (24). We have previously shown that the Ga^3+^MP representative, namely, cationic modified gallium porphyrin (Ga^3+^CHP), is water-soluble and exhibits photodynamic potential at Q-band excitation (at lower absorption peaks) with high antimicrobial activity and low toxicity to human keratinocytes (29,30).

In this study, a human keratinocytes *S. aureus* infection model was constructed and characterized to achieve a stable bacterial presence inside the host cells. We used this model to investigate whether light-activated Ga^3+^MPs could be applied to effectively inactivate extracellular and intracellular *S. aureus*. In our experimental approach, we used three research strategies based on aPDI action (**Fig 1**). Briefly, Strategy 1 refers to the aPDI treatment of *S. aureus* that had escaped from an infected cell **(Fig 1** Strategy 1). Strategy 2 is the aPDI treatment of *S. aureus* cells before they contact and invade keratinocytes (**Fig 1** Strategy 2). Strategy 3 is the aPDI treatment of intracellular *S. aureus* persisting inside host cells (**Fig 1** Strategy 3). In the current study, we used two Ga^3+^MP derivatives that we previously characterized for their antimicrobial efficacy in suspension cells: gallium mesoporphyrin IX (Ga^3+^MPIX) (26) and Ga^3+^CHP (30). We demonstrated that the compounds we studied, in particular Ga^3+^CHP, are able to effectively penetrate host cells, localizing mainly in lysosomal structures, where we also detected the presence of *S. aureus*. Using excitation of Ga^3+^CHP with green light, we achieved a significant reduction in the GFP signal originating from intracellular *S. aureus* by aPDI.

**Fig 1.**
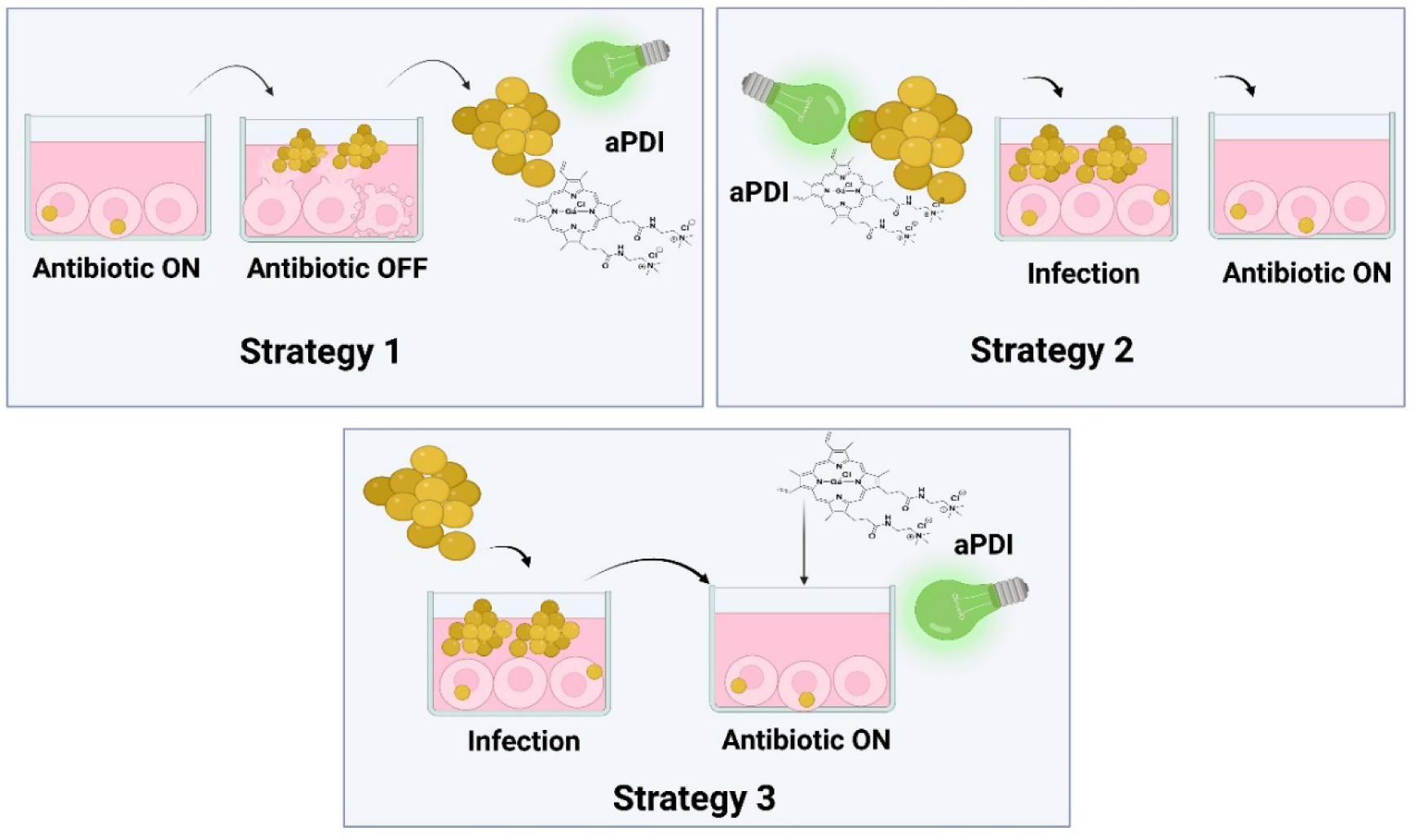
Strategies to implement light-activated gallium metalloporphyrins to overcome *S. aureus* infection of human keratinocytes. Strategy 1 is to reduce the potential for recurrent infection. Strategy 2 investigates how pretreatment with bacterial inoculum before infection affects *S. aureus* adhesion and internalization in host cells. In Strategy 3, photosensitizers are incubated in the dark to efficiently penetrate and localize inside the host cells to reach intracellular *S. aureus*. After incubation, the cells were treated with light.

## Results

### The multiplicity of infection (MOI) affects the infection and internalization of *S. aureus* into human keratinocytes

First, we developed a model of keratinocyte infection (see Materials and Methods, **Fig 11**) and characterized the process of *S. aureus* internalization into keratinocytes. To confirm the presence of intracellular *S. aureus* after infection, fluorescence microscopy images were taken on the first day after infection. The green fluorescent protein (GFP) signal from the *S. aureus* USA300 bacteria (white arrow) was observed inside the host cell along with the nuclei, which were stained with HOEST dye (blue signal, black arrow) (**Fig 2A**). To confirm the intracellular presence of the pathogen, three-dimensional images were taken using scanning fluorescence microscopy (**Fig 2B**), confirming that bacteria were intracellularly localized. Then, we examined the effect of various multiplicities of infection (MOI) (0–100) on the viability of *S. aureus* in medium and inside keratinocytes by studying three fractions after infection. The fractions were as follows: (i) extracellular, (ii) intracellular, and (iii) intracellular + adherent *S. aureus* (**Fig 2C**). The number of extracellular *S. aureus* collected from the culture medium increased with the higher MOI used for infection. For intracellular *S. aureus*, the viability was estimated as 5.6 log_10_ CFU/mL for MOI 100, 4.8 log_10_ for MOI 10, and 4.7 log_10_ for MOI 1 (**Fig 2D)**. By measuring the GFP signal derived from the *S. aureus* strain, a higher number of infected cells was observed for both MOIs 10 and 100 than for an MOI of 1. However, there was no significant difference in the rate of intracellular invasion between the MOI 10 and 100 inoculum according to the flow cytometry studies (**Fig 2E**). The *S. aureus* infection delayed the growth rate of HaCaT cells. However, host cells harboring intracellular *S. aureus* can grow and proliferate. The use of different MOIs was also reflected in host cell growth. The larger the inoculum used for infection, the greater the effect on HaCaT cell growth (**Fig 2F**). The number of viable intracellular pathogens decreased over time as a result of infected cell death (**Movie S1**). Under antibiotic exposure, *S. aureus* USA300 invaded human keratinocytes. The level of *S. aureus* USA300 internalization during this process and the effect of infection on host cell proliferation strongly depend on the initial MOI used for infection.

**Fig 2.**
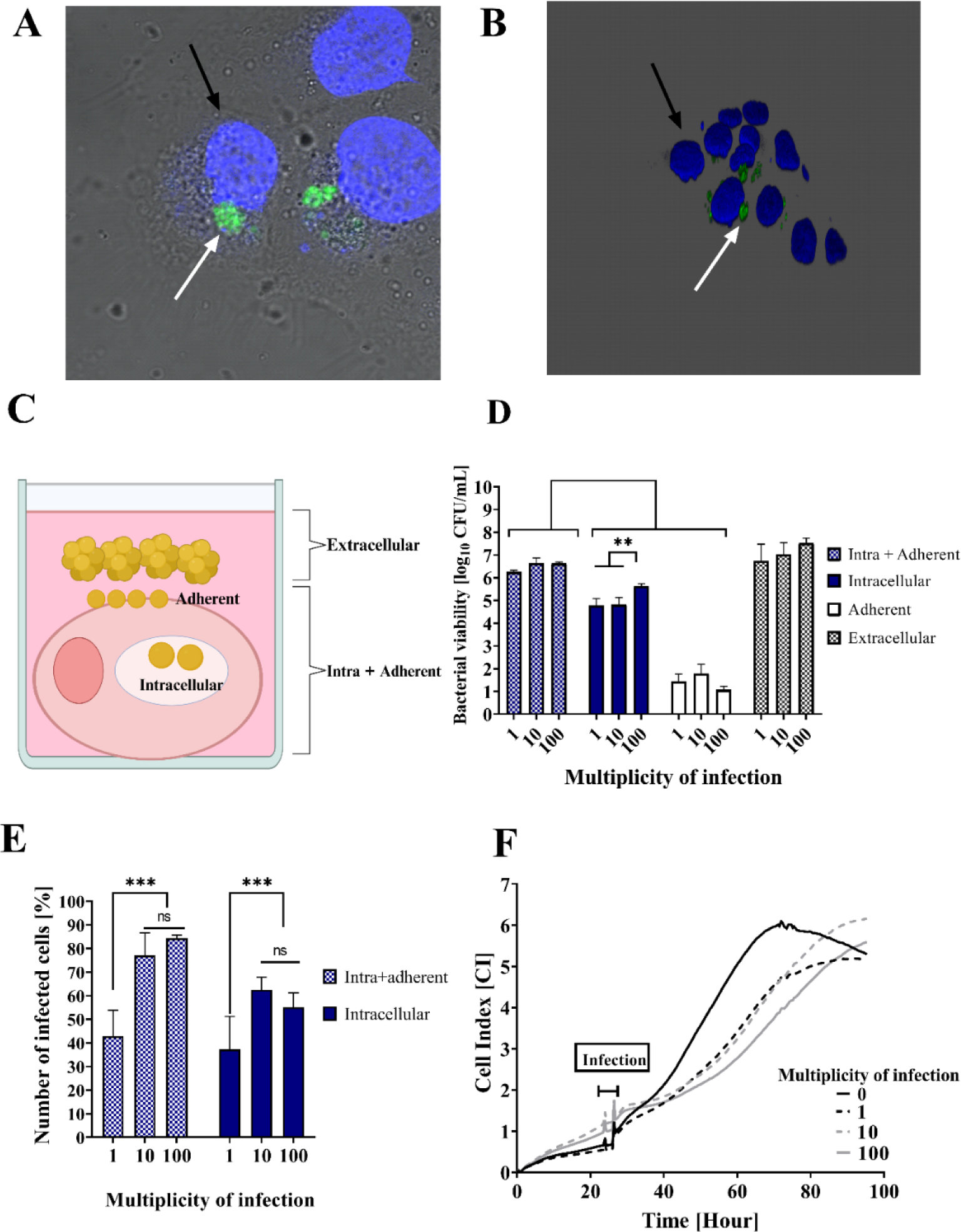
Effect of *S. aureus* multiplicity of infection (MOI) on internalization, infection, and keratinocyte growth. **A)** Keratinocytes (HaCaT cell line) infected with *S. aureus* USA300 (green signal, white arrow). The host cell nucleus was stained (blue signal, black arrow). **(B)** 3D image of the coculture of HaCaT and *S. aureus* USA300. **(C)** Schematic representation of bacterial fractions collected after each step of coculture preparation. ‘Extracellular’ refers to free-floating *S. aureus* collected from the growth medium after a 2-hour staphylococcal infection; ‘Adherent’ - *S. aureus* attached to host cell; ‘Intracellular’ - *S. aureus* accumulated inside host cell; ‘Intra+Adherent’ – the combined number of *S. aureus* in adherent and intracellular fraction. (**D)** Bacterial viability was determined by seeding bacteria onto agar plates and counting colony-forming units (CFU/mL) in each collected fraction. Different ratios of the number of bacteria to the number of host cells were used (MOI 1-100) for infection model preparation. The significant differences are marked with asterisks [**p < 0.01] (**E)** Percentage of infected, GFP-expressing cells collected immediately after infection (MOI 0-100) (‘Intra+Adherent’ fraction) or after 1-hour antibiotic exposure (‘Intracellular’ fraction). The GFP signal was measured by flow cytometry. Significant differences in infected cell viability between tested samples at the respective p values are marked with asterisks [***p < 0.001] and calculated with respect to uninfected cells at each time point. (**F**) Real-time host growth analysis after infection with *S. aureus* USA300 at MOI 0-100. After 2 hours, the medium was removed, the cells were washed, and antibiotics were added to ensure the intracellular maintenance of *S. aureus*.

### The behavior of intracellular *S. aureus* is strain-dependent

To examine whether the staphylococcal invasion into keratinocytes is dependent on the *S. aureus* strain, we compared keratinocyte infections with two *S. aureus* strains: the hypervirulent USA300 strain and the nonvirulent strain RN4220. First, we examined the intracellular viability of the two bacterial strains during infection and their persistence under antibiotic pressure for several days after infection. The nonvirulent RN4220 strain exhibited higher intracellular viability, as shown in a several-day culture model, than the hypervirulent USA300 (5.8 log_10_ vs. 4.2 log_10_ CFU/mL on the 1^st^ day postinfection) (**Fig 3A**). RN4220 remained within the keratinocytes longer than USA300, which was completely titrated out of the cells (reaching the detection limit of 2 log_10_) on day 5 postinfection. The difference in infection between the two strains was also evident in the host growth rate (**Fig 3B**). RN4220 reduced host cell growth after infection significantly more than USA300. Nevertheless, host cells intracellularly harboring bacteria could grow and proliferate (**Fig 3C and D**). Interestingly, when antibiotic pressure was removed and the medium was changed to a nonantibiotic medium (Antibiotic OFF) after the 1^st^ day postinfection (dashed gray line), *S. aureus* RN4220 continued to persisted intracellularly and did not leave the host cells to cause recurrent infection, while USA300 was released from the host, causing cell death, restoring extracellular infection, and inducing significant toxicity. A similar effect was observed when the antibiotic pressure was maintained longer (up to the 3^rd^ day postinfection) for USA300 (**Fig S1**), showing that even the low-viability intracellular inoculum was able to escape and resume the infection. We observed the behavior of *S. aureus* in the infection process, including intracellular persistence and the ability to escape from the host for reinfection, is strain-dependent.

**Fig 3.**
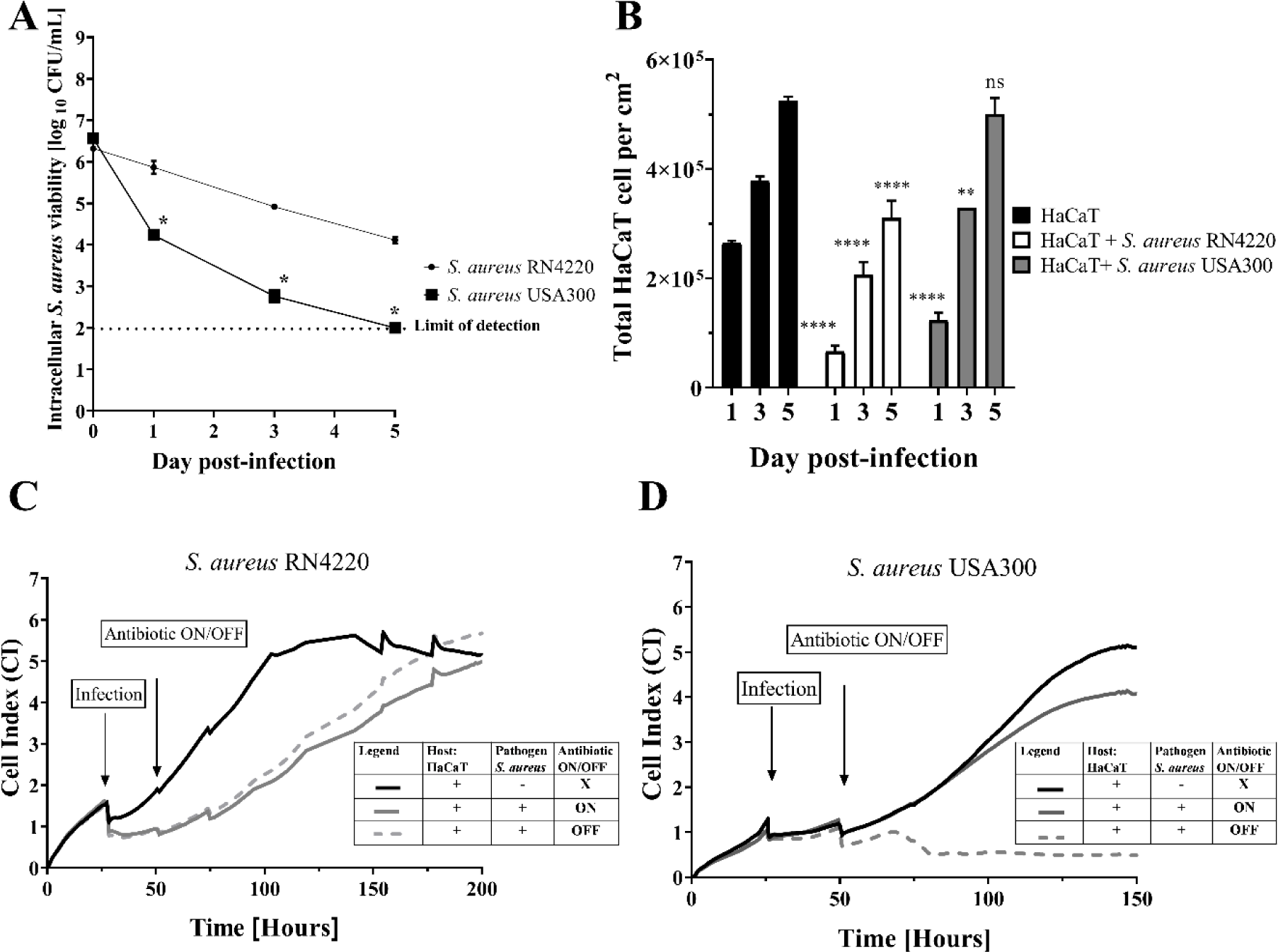
Characterization of intracellular invasion of hypervirulent (USA300) and nonvirulent (RN4220) *S. aureus* strains in an infection model over time. **(A)** Intracellular viability of the two *S. aureus* strains over time after infection (1-5 days). Significant differences in USA300 viability at the tested time points were calculated relative to the reference strain RN4220 with corresponding p values marked with asterisks [*p < 0.05] **(B)** HaCaT cell viability over time (1-5 days) after infection with two tested *S. aureus* strains. The significant differences in infected cell viability at the respective p values are marked with asterisks [*p < 0.05; **p < 0.01; ***p < 0.001] and were normalized using the uninfected cells at each studied time point. Infection of HaCaT cells (**A, B**) in medium without antibiotics was performed at an MOI of 10 for 2 hours; then, the cells were cultured under antibiotic pressure until the end of the experiment. **(C, D)** Real-time analysis of HaCaT cell growth after infection with *S. aureus* RN4220 (**C**) or USA300 (**D**). HaCaT cell infection was performed at an MOI of 10 for 2 hours; no antibiotic (X), culture under antibiotic pressure (Antibiotic ON) or antibiotic removal (Antibiotic OFF), which was determined on day 1 postinfection.

### Filaggrin deficiency promotes the internalization of *S. aureus* and its persistence inside keratinocytes

In AD patients, there is a high mutation frequency of the filaggrin gene (*FLG*), which is linked to increased *S. aureus* skin colonization (31). Filaggrin is a crucial component of the integrity of the epidermal barrier and stabilizes the pH and hydration of the skin to control, for instance, microbial penetration (32). We hypothesized that the presence of filaggrin might be involved in *S. aureus* internalization and persistence in host cells postinfection. To test this hypothesis, two HaCaT cell lines with normal (*FLG* ctrl) and silenced (*FLG* sh) filaggrin status were infected with *S. aureus* USA300. The percentage of infected cells (high expression of GFP) was measured by flow cytometry through days postinfection under constant antibiotic pressure. HaCaT cells with silenced filaggrin expression (*FLG* sh) showed a significantly higher number of infected cells after infection than cells with normal filaggrin expression (83%±3.8 vs. 71%±4.3) (**Fig 4A**). This tendency continued until the 5^th^ day postinfection under antibiotic pressure. Intracellular *S. aureus* persists longer inside keratinocytes with filaggrin dysfunction (up to 7 days). Moreover, we observed delayed growth of the filaggrin-silenced cell line relative to the wild-type line due to *S. aureus* infection (**Fig 4B**). However, both cell lines were still able to proliferate with continuous clearance of intracellular infection. Intracellular *S. aureus* was gradually titrated out of the cells that were able to reach the plateau phase (*FLG* ctrl cell line at 55 hours of experiment and *FLG* sh cell line at 85 h). Impairment of functional filaggrin in the host cell may be promote the internalization of *S. aureus* and its intracellular persistence.

**Fig 4.**
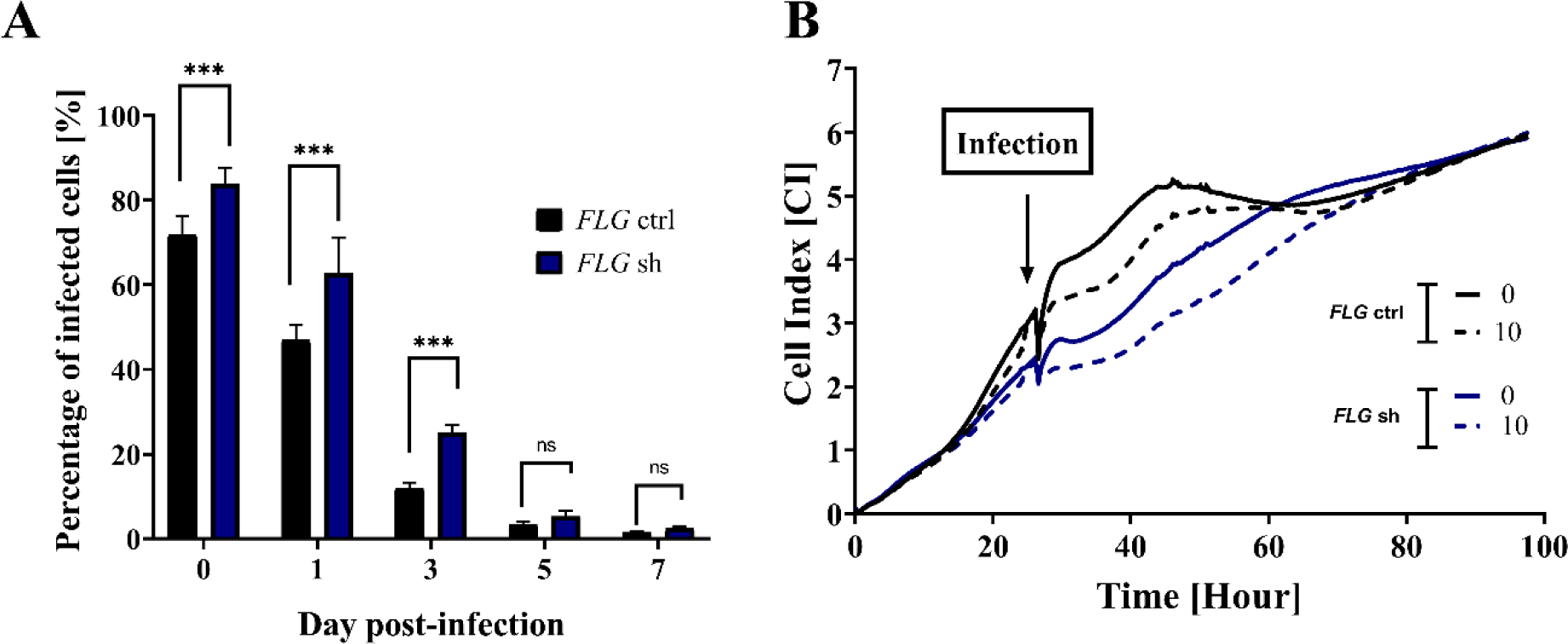
Filaggrin status affects the staphylococcal invasion into human keratinocytes. **A)** Percentage of GFP-expressing infected cells in culture with divergent filaggrin (*FLG*) expression. HaCaT cells with normal expression of filaggrin (*FLG* ctrl) and silenced filaggrin expression (*FLG* sh) were infected with *S. aureus* USA300 at an MOI of 10 for 2 hours. Antibiotic pressure was then maintained for up to 7 days. Cells were harvested at each time point after infection, and the number of GFP-expressing cells was determined by flow cytometry. Significant differences are marked with asterisks [***p < 0.001] and were calculated with respect to *FLG* ctrl cells at each time point. **B)** Real-time growth analysis of both cell lines infected and noninfected with staphylococcal cells with an MOI of 10 for 2 hours. After the initial infection, antibiotic pressure was maintained throughout the whole analysis.

### Green light-activated Ga^3+^MPs efficiently eliminate recurrent staphylococcal infection

Our previous study showed that light-activated Ga^3+^MPs are able to reduce the viability of clinical *S. aureus* isolates in suspension cultures *in vitro* (29). In the current study, we wanted to verify whether and at what stage of the infection cycle light-activated Ga^3+^MPs (i.e., aPDI) can be used to reduce the number of bacteria that infect host cells. Based on the first proposed aPDI strategy (**Fig 1, Strategy 1**), we evaluated the antimicrobial efficacy of Ga^3+^MPs activated with 522 nm light against *S. aureus* USA300 in staphylococcal reinfections (**Fig 3D**, dashed line). Briefly, cells were seeded on day 0, *S. aureus* infection was performed on day 1, and then cells were cultivated in medium with an antibiotic (Antibiotic ON) (**Fig 5A**). On day 2, the culture medium was changed to antibiotic-free medium (Antibiotic OFF), and the cells were cultivated for up to 16 hours. Extracellular bacteria that were released from keratinocytes were isolated, washed, and subjected to aPDI using two Ga^3+^MPs, Ga^3+^MPIX or Ga^3+^CHP, to assess their efficacy in reducing *S. aureus* survival rates (**Fig 5B and 5C**). Both gallium compounds effectively eliminated bacteria released from cells with a maximum reduction in cell count of 4 log_10_ CFU/mL. Ga^3+^CHP was more effective than Ga^3+^MPIX in reducing bacterial survival at lower light doses, corresponding to shorter irradiation times. We found that *S. aureus*, which was released by the host and restarted the infection, responded to aPDI like bacteria in the logarithmic growth phase rather than the stationary phase (**Fig S2**). Despite some adaptive changes in intracellular bacteria that might promote their overall tolerance, the released *S. aureus* showed a similar response to aPDI as that of bacteria in free suspension cultures during the logarithmic growth phase. This finding demonstrated that aPDI with Ga^3+^MPs can be an effective strategy to combat staphylococcal reinfections.

**Figure 5.**
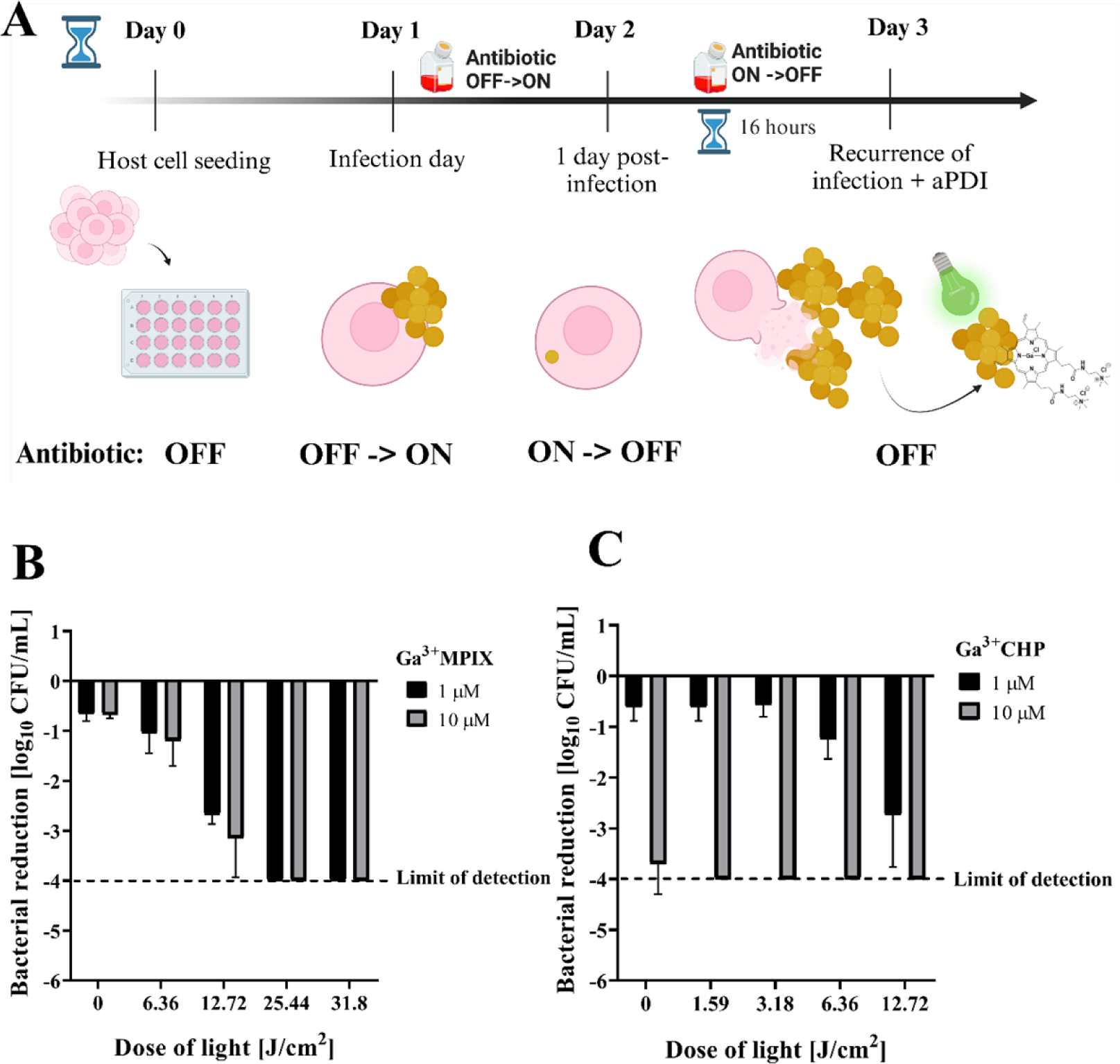
Light-activated Ga^3+^MPs effectively reduce the number of *S. aureus* USA300 released from keratinocytes – Strategy 1. (A) On day 1, HaCaT cells were infected with *S. aureus* USA300 (MOI = 10, 2 hours) in medium without antibiotics (Antibiotic OFF). Antibiotic was added to remove extracellular *S. aureus,* and the cells were cultured under antibiotic pressure until the next day (Antibiotic OFF->ON). The antibiotic was then removed (Antibiotic ON->OFF), and HaCaT cells containing only intracellular *S. aureus* were left in an incubator for 16 hours. During this time, intracellular *S. aureus* was gradually lysed cells and released into the medium. *S. aureus* cells were harvested and washed with DMEM and then resuspended in tryptic soy broth (TSB). Bacterial suspensions were transferred to a 24-well plate, proper photosensitizers were added, and after incubation, they were illuminated with green light. Bacterial cells were then diluted, plated, and counted (B and C). The reduction in the number of *S. aureus* USA300 bacteria by light activation of Ga^3+^CHP (B) or Ga^3+^MPIX (C) was calculated in relation to the untreated cells.

### aPDI reduces the severity of *S. aureus* infection and its adherence to the host cell

We hypothesized that aPDI pretreatment of the infectious inoculum could affect its further invasion, adherence to host, and growth of the extracellular bacteria (**Fig 1, Strategy 2**). To assess the effect of aPDI pretreatment on the initial stages of *S. aureus* invasion, we studied the number of bacteria in each fraction collected after infection. As a control, we used nontreated infectious inoculum (**Fig 6**). Notably, we used the same number of bacteria regardless of the treatment. Furthermore, we wanted to investigate the severity of the infection, so two MOI values were used, 10 and 1 to investigate higher and lower amounts of bacteria. For aPDI-treated bacteria, two doses were used: low and high to reduce bacterial viability to an appropriate number of bacteria. The low dose was used to obtain a higher number of bacteria at an MOI of 10 (**Fig 6B**, Low), while the high dose was used to obtain a lower number of bacteria at an MOI of 1 (**Fig 6B**, High). After infection, the CFU/mL were counted in (i) the extracellular fraction from the growth medium after infection, (ii) the intracellular+adherent fraction obtained from a cell lysate containing both intracellular and adherent bacteria, or (iii) the intracellular fraction from a cell lysate where HaCaT cells were washed and cultured under antibiotic pressure for 1 h before lysis to eliminate adherent bacteria (see **Fig 6A**). The aPDI-treated bacteria used as inoculum for infection behaved differently from the untreated bacteria. Light-activated Ga^3+^CHP treatment of *S. aureus* resulted in a significantly reduced number of extracellular bacteria, and the observed decrease was by 2 log_10_ for low-dose treatment and 2.8 log_10_ for high-dose treatment compared to untreated controls (**Fig 6B**). Moreover, high-dose aPDI resulted in a significant reduction in bacterial adherence to HaCaT cells, with a 1.2 log_10_ reduction in CFU/mL count compared with the adherence of untreated bacteria. Notably, both aPDI-treated bacteria and nontreated bacteria were applied at the same MOI of 1, indicating that the quality of bacteria but not their quantity influenced adherence. Interestingly, pretreatment of bacteria with aPDI did not affect the number of intracellular bacteria, suggesting that there is a maximum yield of bacterial burden that can invade host cells, and it is not affected by aPDI. Both aPDI-treated and untreated bacteria penetrated keratinocytes with similar efficiency. However, aPDI treatment itself significantly affects the growth of the extracellular fraction and bacterial adhesion to host cells.

**Figure 6.**
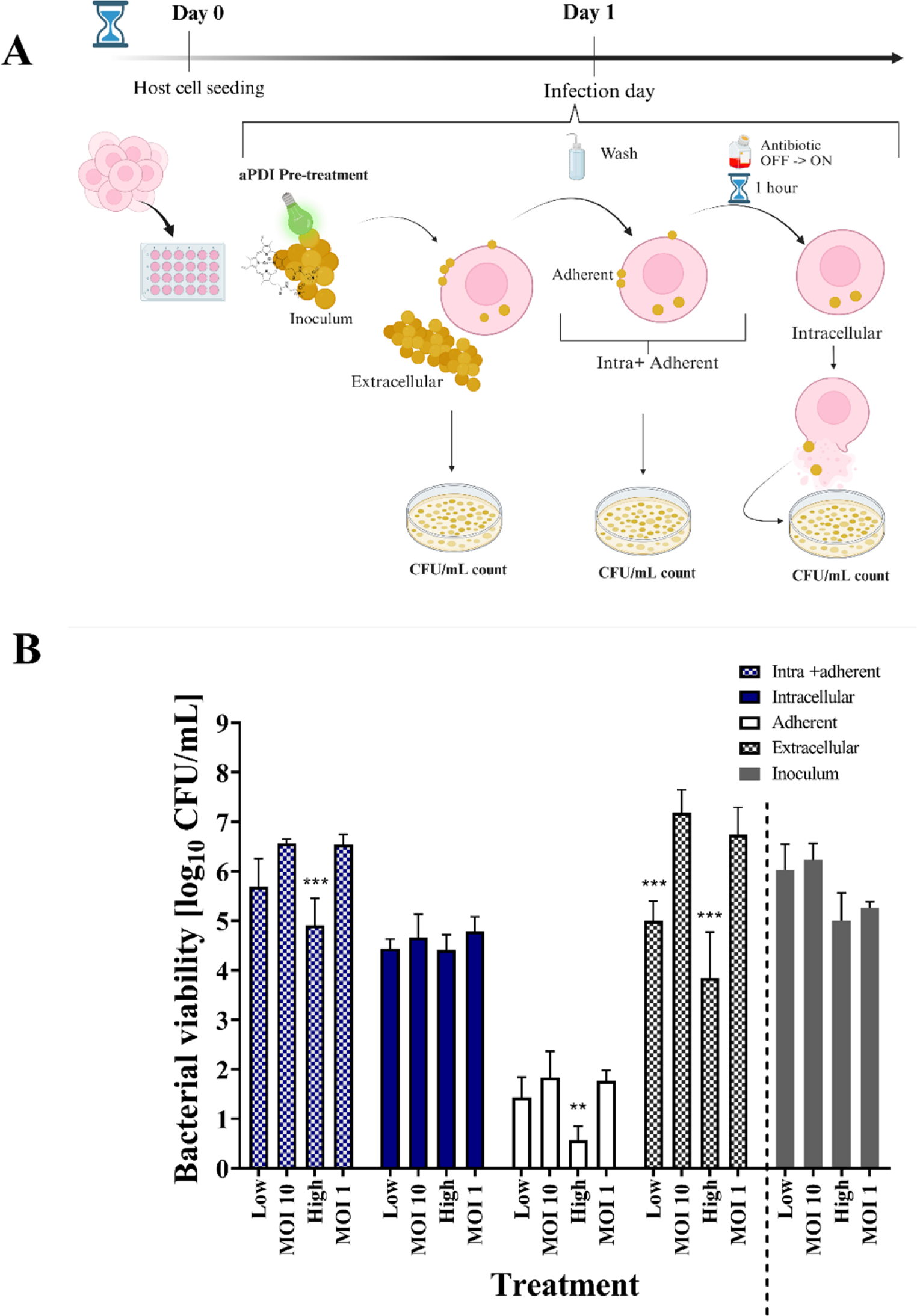
Effect of aPDI (light-activated Ga^3+^CHP) pretreatment on *S. aureus* USA300 infection – Strategy 2. **(A)** On day 0, HaCaT cells were seeded in a 24-well plate. Then, on day 1, *S. aureus* USA300 infection was conducted either with nontreated bacteria or bacteria treated with aPDI. After 2 h, the growth medium was collected for plating and counting the extracellular fraction. Host cells were collected for subsequent lysis (intra+adherent fraction) or incubated for 1 hour with antibiotics to eliminate adherent bacteria to obtain the intracellular fraction. All fractions were serially diluted and plated for CFU/mL enumeration. **(B)** The number of bacteria counted from each fraction collected after infection with untreated or aPDI-treated *S. aureus* inoculum. Before infection, the number of viable *S. aureus* (10^7^ CFU/mL) was reduced by two doses of aPDI: a low dose that resulted in a 1 log_10_ reduction in CFU/mL (MOI 10) and high dose that resulted in a 2 log_10_ reduction in CFU/mL (MOI 1). Untreated bacteria with the appropriate MOI of 10 or 1 were used as a control. The number of bacteria used for infection was the same whether the bacteria were pretreated with aPDI or were untreated (please see Inoculum). After infection, several fractions were collected, such as Extracellular – the free-floating *S. aureus* collected from the medium after infection; Adherent - *S. aureus* attached to the host cell; Intracellular - *S. aureus* accumulated inside the host cell in the presence of antibiotic pressure; Intra+Adherent – combined number of intracellular and adherent *S. aureus*; Inoculum – initial number of treated (Low or High) or untreated (MOI 10 or MOI 1) *S. aureus* used for infection. The data are presented as the mean ± SD of six separate experiments. The significance differences in infected cell viabilities at the respective p values are indicated with asterisks [**p < 0.01; ***p < 0.001], and were normalized to the respective control of each pretreatment.

### Variable patterns of Ga3+ MP accumulation inside human keratinocytes

In the next stage of our work, it was important to carefully examine the accumulation of the tested compounds in host cells before applying Strategy 3 (**Fig 1** Strategy 3), which involves applying aPDI to intracellular bacteria. First, we investigated the accumulation profile of gallium compounds, Ga^3+^CHP and Ga^3+^MPIX inside human keratinocytes to check for possible differences and the efficacy of the process. The accumulation of Ga^3+^ MPs is a crucial step for reaching the bacteria inside keratinocytes. The rate of accumulation of both compounds was time-dependent. After 1 hour of accumulation, both Ga^3+^CHP and Ga^3+^MPIX were primarily localized at the cell membrane (**Fig 7A**). However, over time, we observed differences in the localization patterns of the tested gallium derivatives. Ga^3+^MPIX was distributed throughout the cell in the cytoplasm, while cationic Ga^3+^CHP was mainly localized inside the cell in clusters (**Fig 7A**). Flow cytometry analysis revealed that after 2 hours of accumulation, 60% of keratinocytes accumulated Ga^3+^MPIX, while the corresponding value for Ga^3+^CHP was 7% (**Fig 7B**). After 6 hours, 75% of the cells accumulated Ga^3+^MPIX and 59% Ga^3+^CHP, reducing the differences observed at the 1st and 2nd hours. Interestingly, after 24 hours, nearly every cell accumulated both compounds at the same level (94%). When measuring the Ga^3+^CHP or Ga^3+^MPIX accumulation in each cell, we also observed time-dependent uptake (**Fig 7C**). Remarkably, despite the accumulation in the cells, none of the tested compounds exerted a significant cytotoxic effect against keratinocytes (**Fig 7D, E**). Even after a 24-hour incubation with the tested compounds, the cells continued to grow and proliferate until finally reaching the plateau phase at a similar time, regardless of the incubation time with the compound. Based on these results, both compounds accumulate in host cells in a time-dependent manner without significantly affecting host proliferation and viability. However, the accumulation patterns of Ga^3+^MPIX and Ga^3+^CHP inside the cell are highly divergent. It is more likely for Ga^3+^CHP, than Ga^3+^MPIX, to reach intracellular *S. aureus* due to its more localized accumulation inside the cells.

**Figure 7.**
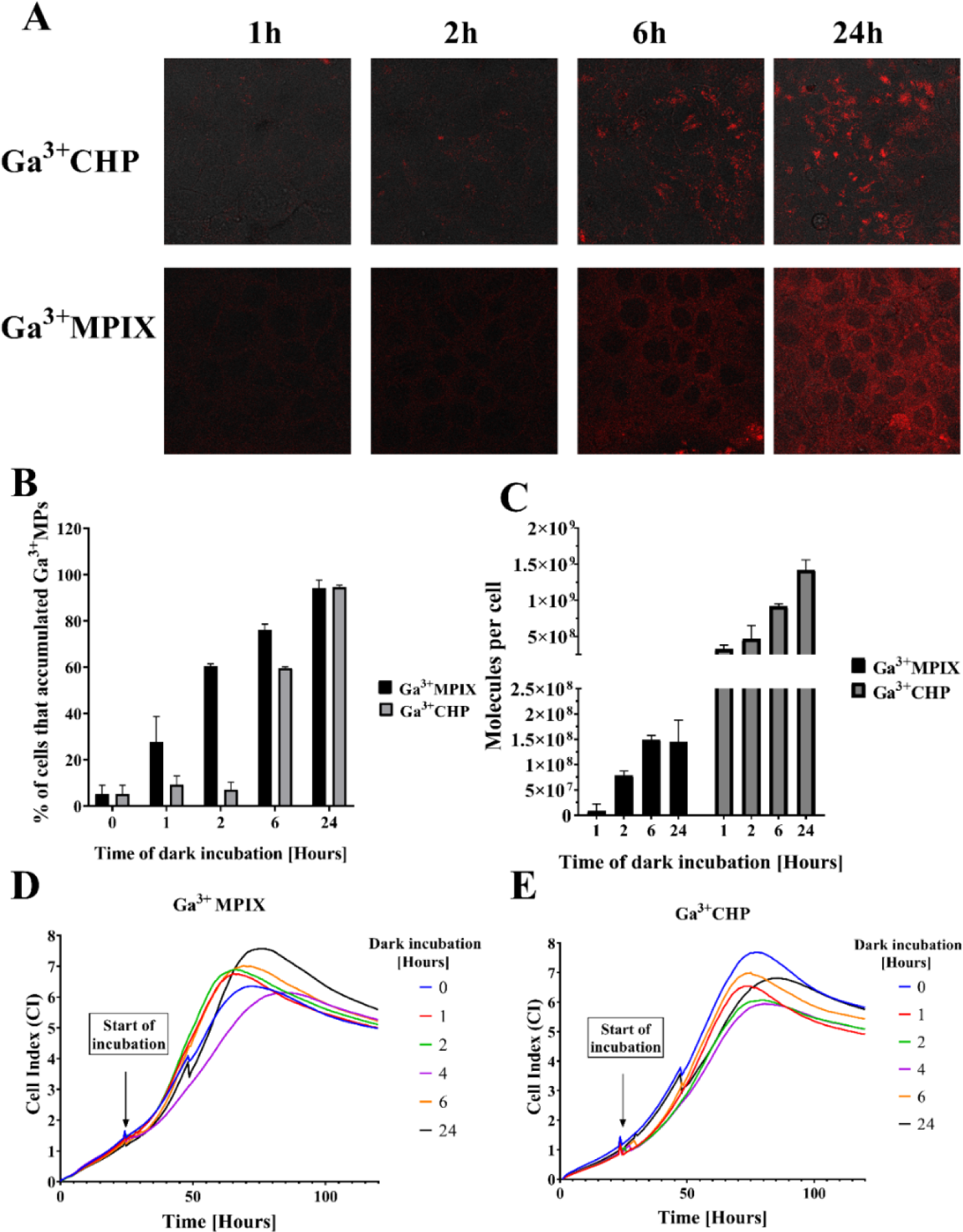
Different accumulation patterns between two gallium metalloporphyrins in human keratinocytes. (**A**) Confocal fluorescent microscopy images showing the accumulation of two gallium compounds, Ga^3+^MPIX and Ga^3+^CHP, in HaCaT cells after incubation in the dark (1-24 hours) at a 10 µM concentration. (**B**) Percentage of cells that accumulated Ga^3+^MPs among the total number of cells measured by flow cytometry. Cells were incubated with 10 µM Ga^3+^MPIX or Ga^3+^CHP and fixed at each time point (1-24 hrs) in the absence of light, and then the fluorescence signal in the cells was measured by flow cytometry. (**C**) The number of accumulated Ga^3+^MPIX or Ga^3+^CHP molecules (10 µM) in keratinocytes after incubation for the time indicated on the X axis, as measured by the fluorescence intensity of the cell lysate. After incubation in the dark for a particular time, the cells were harvested, counted, and lysed with 0.1 M NaOH/1% SDS to determine the fluorescence of each accumulated compound. (**D, E**) Real-time growth analysis of HaCaT cells after dark incubation (1-24 hours) with 10 µM Ga^3+^MPIX (D) or Ga^3+^CHP (E).

### The accumulation of Ga^3+^CHP was greater in infected cells

We next investigated the accumulation of Ga^3+^CHP in *S. aureus*-infected cells. First, keratinocytes were infected with *S. aureus* for 2 hours (MOI 10) in medium without antibiotics, and then antibiotics were introduced to remove extracellular bacteria and maintain intracellular invasion. For comparison, uninfected cells were separately cultured. The next day, Ga^3+^CHP was added to the infected cell culture, and the cells were incubated in the dark. After 2, 4, and 6 hours of incubation, cells were collected to detect the red fluorescence signal from the accumulated compound in both infected and uninfected cells (**Fig 8A**). We observed an increase in the accumulation level over time, measured as the percentage of keratinocytes (infected or noninfected) showing fluorescence produced by Ga^3+^CHP. We observed a very interesting correlation in that infected keratinocytes accumulated more Ga^3+^CHP than uninfected cells. This observation was most evident after a longer incubation (**Fig 8A**, 6 hours). Then, we analyzed the compound’s accumulation only in the fraction of infected cells containing both signals: red fluorescence from the cationic Ga^3+^CHP compound (Ga^3+^CHP+) and green fluorescence from the *S. aureus* strain USA300 (GFP+) (**Fig 8B**). Over time, the accumulation of a gallium compound in the infected cell fraction (Ga^3+^CHP+/GFP+) increased. After 6 hours of incubation, the number of infected cells simultaneously possessing both signals increased up to 25%, indicating that the cationic gallium compound accumulates in infected cells. This finding indicated that cationic Ga^3+^CHP may colocalize with intracellular bacteria.

**Figure 8.**
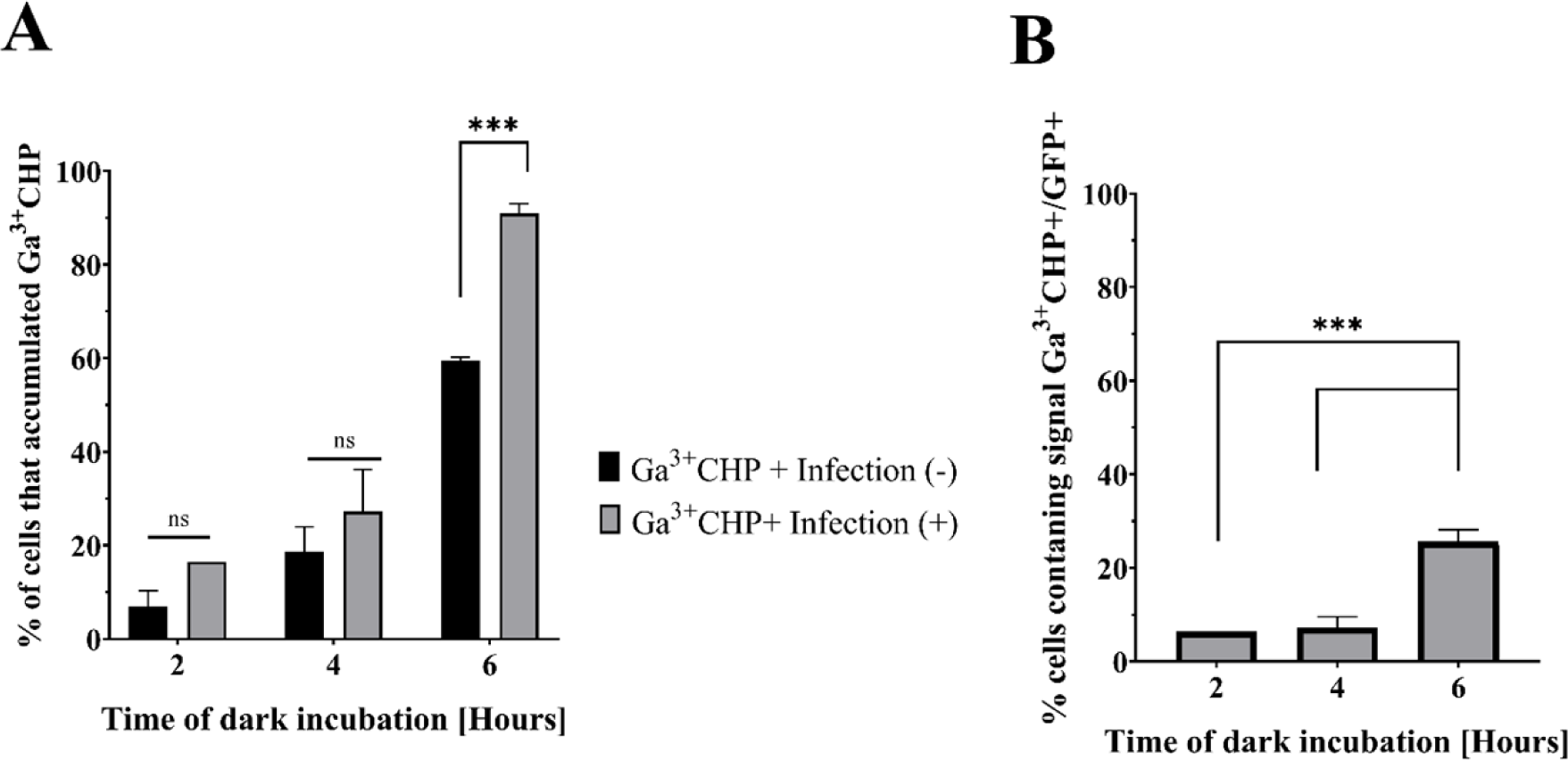
The accumulation of Ga^3+^CHP in infected cells. (A) Accumulation of Ga^3+^CHP in infected and uninfected cells. The cells were infected with *S. aureus* USA300 (MOI 10) for 2 hours in medium without antibiotics, and then the cells were washed and covered with medium with antibiotics. In parallel, uninfected cells were cultured. The next day, 10 µM Ga^3+^CHP was added and incubated in the dark. After incubation, the cells were washed and collected, and the percentage of cells with a fluorescence signal derived from Ga^3+^CHP was determined by flow cytometry. The accumulation results were compared with those of uninfected cells. (B) The number of cells harboring both GFP+ and Ga^3+^CHP+ signals after incubation in the dark with Ga^3+^CHP in infected cells. The significant differences between the two compounds were calculated, and the respective p values are marked with asterisks [***p < 0.001].

### Intracellular *S. aureus* colocalizes with Ga^3+^CHP in keratinocyte lysosomes

Ga^3+^CHP accumulated inside the cells in distinct clusters; thus, in the next experiment, we wanted to identify the organelles in which the studied compound localized. Fluorescence dyes specific for the Golgi apparatus, mitochondria, and lysosome were used to determine the localization of the compound (**Fig S3, Table S1**). To assess colocalization, images were analyzed for two colocalization coefficients: the Pearson correlation (significant colocalization > 0.5) and overlap coefficient (> 0.6). High localization of the compound in lysosomes was observed, but this was not the only site of accumulation. The overlap coefficient between Ga^3+^CHP and LysoTracker™ Deep Red in lysosomes was 0.65 with a Pearson’s correlation of 0.46 (**Fig 9**), indicating partial localization in lysosomes. We also observed some Ga^3+^CHP accumulation in the mitochondria (although to a far lesser extent compared with lysosomes) and no confirmed localization in the Golgi apparatus (**Table S1**). Then, we examined the localization of *S. aureus* inside keratinocytes. However, we did not confirm the lysosomal localization of the bacteria in keratinocytes by both the overlap coefficient and Pearson’s correlation (**Table S2**). Interestingly, when Ga^3+^CHP was incubated with *S. aureus*-infected keratinocytes, colocalization between *S. aureus* and Ga^3+^CHP occurred. Moreover, simultaneous staining of lysosomes and detection of signals from *S. aureus* (GFP) and Ga^3+^CHP indicated the colocalization of these signals in the lysosomes. We thus hypothesized that certain metabolic changes caused by two factors (infection and presence of Ga^3+^CHP) can promote the colocalization of *S. aureus* inside lysosomes.

**Figure 9.**
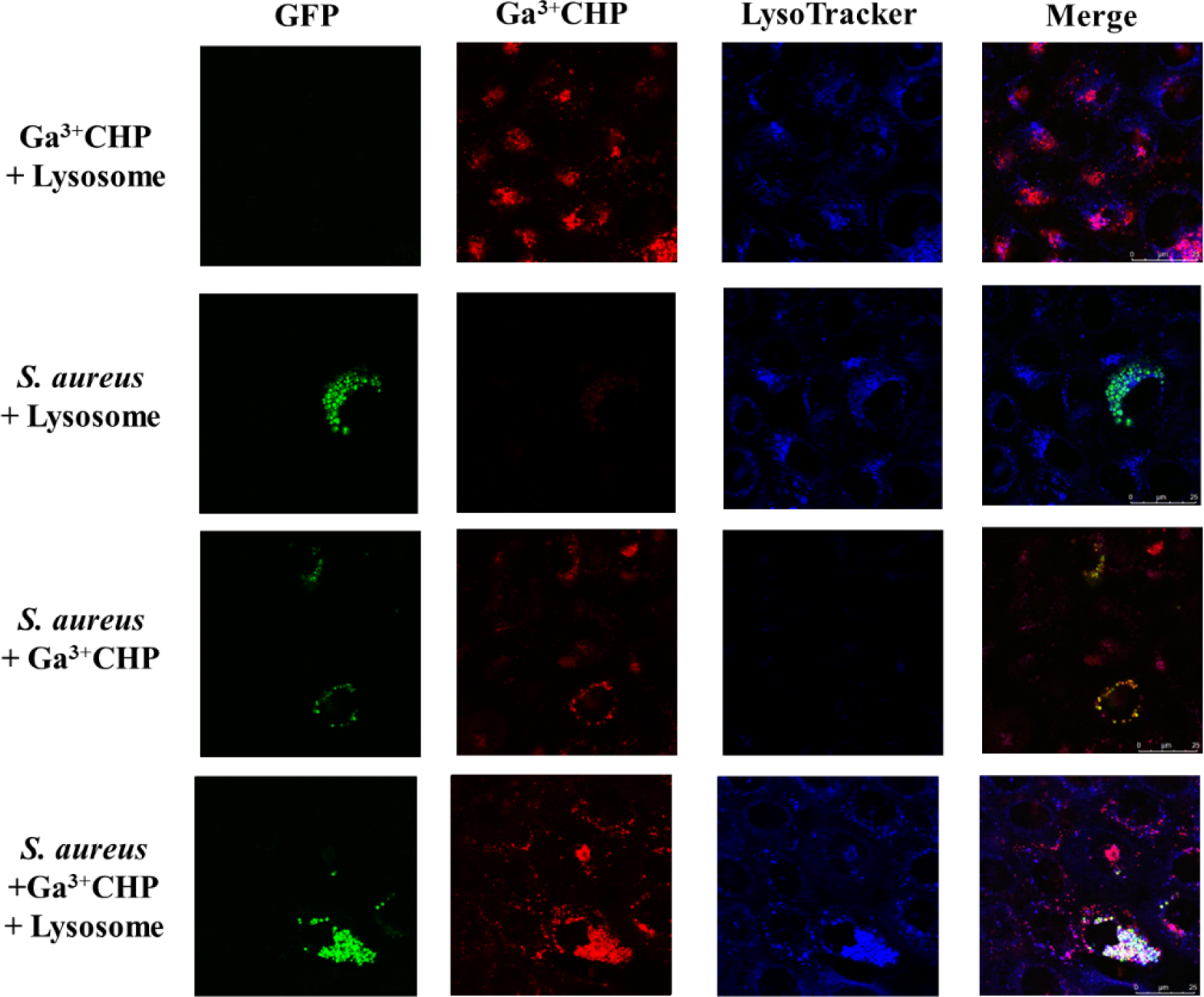
Colocalization of intracellular *S. aureus* and Ga^3+^CHP in lysosomes. Confocal images of *S. aureus*-infected keratinocytes (HaCaT) on the 1^st^ day postinfection, incubated in the dark for 6 hours with cationic Ga^3+^CHP (10 µM). The green signal represents intracellular *S. aureus* USA300-GFP, the red signal is Ga^3+^CHP, and the blue signal represents lysosomes. All colocalization parameters are detailed in **Table S2** in the Supporting Information Captions.

### Light-activated Ga^3+^CHP reduces the number of infected keratinocytes

After establishing the colocalization of intracellular bacteria and Ga^3+^CHP, we proceeded to analyze the effect of aPDI on intracellular and intralysosomal *S. aureus* according to Strategy 3 (**Fig 1** Strategy 3). To this end, we excited the accumulated Ga^3+^CHP in the cells with green light and investigated how it affected the total number of infected cells. For this purpose, on the first day, we infected keratinocytes, removed extracellular bacteria, and left only intracellular bacteria by adding antibiotics in the culture. The next day, we added Ga^3+^CHP to the cells and incubated them for 2 or 6 hours in the dark. After incubation, we irradiated the cells with green light at doses of 6.36 or 12.72 J/cm^2^ (**Fig 10A**). Using flow cytometry, we determined the number of infected cells by tracing the GFP signal originating from intracellular *S. aureus*. As a control (100%), the GFP signal of infected cells that were not treated subjected to aPDI was analyzed. Incubation of cells with Ga^3+^CHP in the dark reduced the number of cells expressing GFP to ∼70%, regardless of incubation time. However, when green light irradiation was added, a significant decrease in the number of GFP-expressing cells to 37-27% was observed, which was not dependent on incubation time or the light dose applied (**Fig 10B**). Treatment with light alone did not reduce the number of infected cells. For Ga^3+^MPIX, which did not accumulate in intracellular clusters, we did not observe any decrease in the number of infected cells after green light illumination (**Fig S4**). Colocalization of both the photosensitizer and the bacteria is crucial for the effective reduction of the intracellular *S. aureus* during the aPDI process.

**Figure 10.**
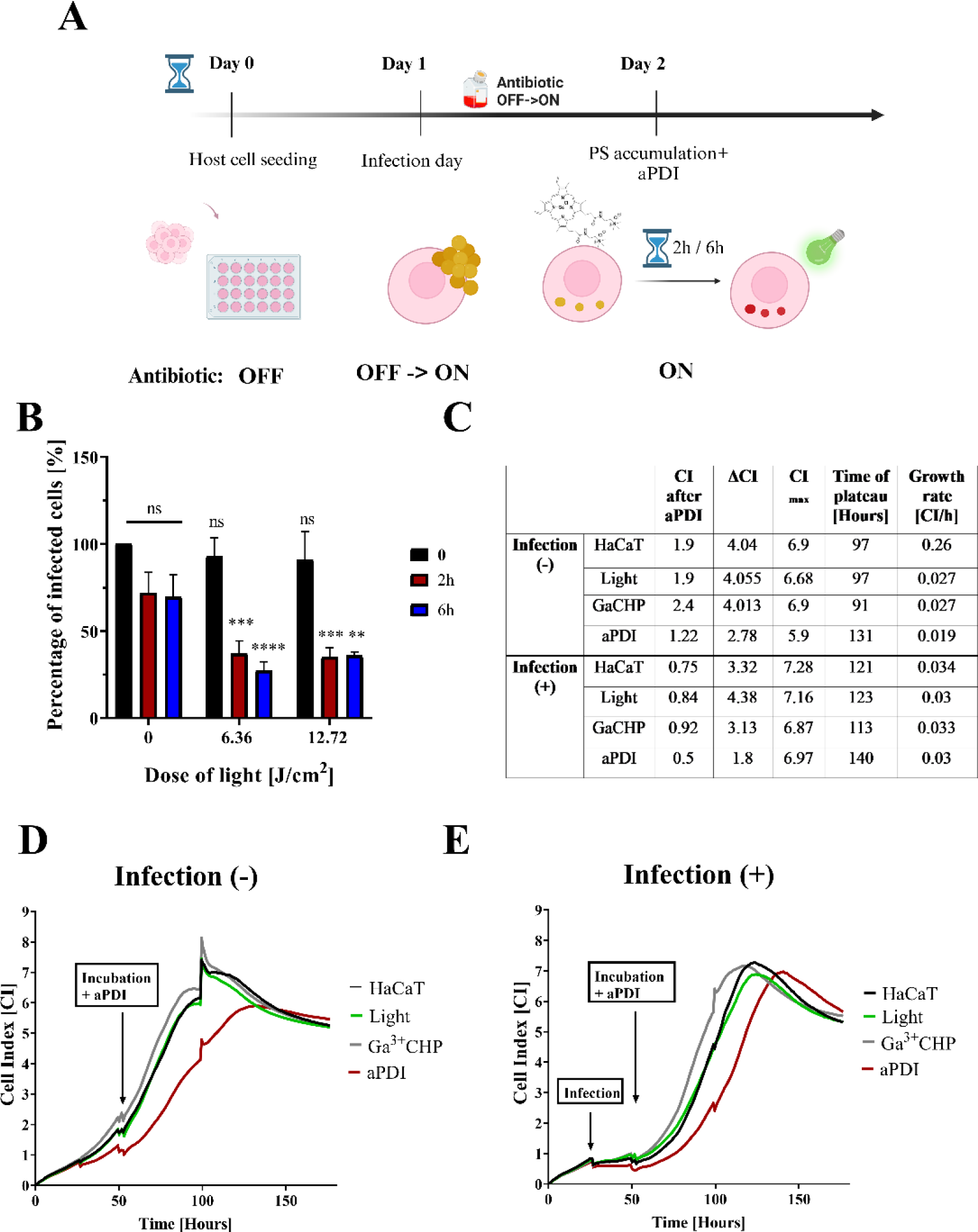
Light-activated Ga^3+^CHP impacts the number of GFP-expressing cells without causing severe host damage. (A) Scheme of the experiment. On day 0, HaCaT cells were seeded in a 24-well plate. On day 1, *S. aureus* USA300 (MOI 10) was added and incubated for 2 hours. Next, the medium was replaced with antibiotic medium (Antibiotic OFF->ON) to eliminate extracellular bacteria and maintain intracellular invasion. On day 2, Ga^3+^CHP was added to the medium and incubated for 2 or 6 hours in the dark. Then, the cells were washed, and green light was applied at the proper dose. (B) Percentage of infected cells after aPDI. The number of GFP-expressing cells after 2-or 6-hour incubation in the dark followed by green light illumination. Cells were collected and fixed, and the GFP signal was measured by flow cytometry. The results were calculated in reference to the untreated control (cells with no compound and no light exposure). (C) Table containing the growth characteristics of infected or uninfected cells administered different treatments. The following parameters were accounted for in the analysis: the cell index (CI) immediately after aPDI treatment (at the 55^th^ hour of the experiment); ΔCI with the characteristics of the growth rate immediately after aPDI treatment, calculated as the difference in the growth of cells after aPDI treatment (from 55 h of the experiment) and cells in the middle of logarithmic growth (∼100 h of the experiment); CI_max_-Maximum CI achieved at the beginning of the plateau phase; time to plateau phase - time at which cells enter stationary growth phase; growth rate calculated as the total rate of increase in the logarithmic phase of growth after treatment to time of reaching the plateau phase of the analysis curve. (D, E) Real-time growth analysis of HaCaT cells uninfected (D) or infected (E) after incubation in the dark for 2 hours with 10 µM Ga^3+^CHP and with or without light illumination of 6.36 J/cm^2^. The CI was measured by a real-time cell analysis (RTCA) instrument every 10 minutes. Experiments were conducted until the cells reached the plateau growth phase.

To verify how aPDI treatment affects the survival and growth of host cells, we analyzed the real-time growth of infected and noninfected cells after illumination with green light (6.36 J/cm^2^) following a 2-hour incubation with Ga^3+^CHP in the dark (**Fig 10 C-E**). We observed that incubation of cells with Ga^3+^CHP followed by light application delayed the proliferation rate of both infected and uninfected cells. Interestingly, after aPDI, infected cells exhibited a higher overall growth rate (0.03 CI/h) than uninfected cells (0.019 CI/h) until reaching the plateau phase. However, cell growth resumed immediately after aPDI in uninfected cells compared to infected cells (as reflected by the ΔCI parameter: 2.78 vs. 1.8 for noninfected vs. infected cells, respectively). The plateau phase was reached faster in uninfected cells (131 h, CI_max_ = 5.9) than infected cells (140 h, CI_max_ = 6.9). Compared to untreated cells, both infected and uninfected cells showed a significant reduction in growth and a delay in entering the plateau phase after aPDI treatment. Despite some phototoxicity of aPDI, which reduces the rate of cell proliferation after treatment, both infected and uninfected cells overcame photodamage and began to proliferate until the plateau phase is reached. This finding indicated a lack of significant photo-and cytotoxicity against host cells.

## Discussion

In patients with chronic and recurrent skin infections, such as AD, preventing and treating staphylococcal infections are extremely important. The interactions between *S. aureus* and skin cells are the subject of intensive research. On the one hand, investigations on whether there are specific characteristics that allow bacteria to adapt to living on the skin are being conducted. On the other hand, the host cells are the research focus, which are not just passive players in the host‒pathogen interaction. Methods are constantly being developed to control the spread of staphylococci on skin, which happens frequently in individuals with skin diseases, such as AD. Antibiotics are used to combat staphylococci skin infection, but the frequent use of antibiotics leads to the selection of strains resistant to the drug used. In this case, other methods are needed, preferably those that can combat infections caused by drug-resistant bacteria. One method that is described in this article is aPDI. It is a method with proven efficacy against not only *S. aureus* but also other microorganisms, including drug-resistant pathogens (21). However, it has not yet been shown whether the use of aPDI is effective against intracellular bacteria.

*Staphylococcus aureus* was initially described as an extracellular pathogen, but recent scientific reports indicate its ability to invade nonprofessional phagocytes, including keratinocytes. In this way, the bacteria avoid the effect of antibiotics, which cannot penetrate the host cell membrane (6,33). In our study, we established a keratinocyte infection model and investigated the efficacy of aPDI in treating multidrug-resistant *S. aureus* at the different infection stages: adherence, internalization, intracellular persistence, or release of bacteria from the host cell (**Fig 1**). In this study, we used the HaCaT cell line and keratinocytes with reduced filaggrin function as a simplified model of AD. The results of our study presented in this work remain in agreement with previously published data showing that in patients with AD, the impairment in the *FLG* gene is correlated with increased *S. aureus* colonization (32,34). We observed a higher internalization and longer intracellular persistence of *S. aureus* in keratinocytes without functional filaggrin (**Fig 4**). Higher intracellular *S. aureus* persistence might be responsible for chronic staphylococcal infections in AD patients and an increased risk of reinfection after initial antibiotic treatment (35).

Once internalized, *S. aureus* is exposed to two selective pressures: antibiotics and the intracellular environment of the host. The pathogen undergoes drastic transcriptional changes to enhance persistance (36,37). Despite these adaptive changes, intracellular bacteria still exhibit metabolic activity. Moreover, it has been shown that the intracellular phenotype could manifest greater tolerance to antibiotics (38). The reservoir of intracellular *S. aureus* is strongly linked to the recurrence of infection (39–41). Under favorable conditions (i.e., the absence of antibiotics), *S. aureus* USA300 could escape the cells and reinduce the infection, even with a lower intracellular load (**Fig 3D, Fig S1**). The same observation was made by Rollin et al., where *S. aureus* resumed extracellular infection after maintaining intracellular inoculum for up to 10 days under selective antibiotic pressure (42). Bacteria that are released from the host might contribute to therapeutic failures. In our study, we evaluated the efficacy of aPDI action (Strategy 1) in eliminating *S. aureus* released from host cells that had regrown from the intracellular inoculum. Based on our results, bacteria released from cells were still sensitive to aPDI, similar to suspension cultures of *S. aureus* grown *in vitro* (**Fig 5**). The adaptive changes that occurred during infection did not alter the bacterial sensitivity to aPDI. These findings indicate that the combination of green light and Ga^3+^MPs might be an efficient strategy to combat recurrent staphylococcal infections.

In Strategy 2, we assessed the effect of aPDI pretreatment on *S. aureus* adherence and internalization. Treating the initial inoculum with Ga^3+^CHP before infection significantly decreased the adherence of bacteria to keratinocytes. In our previous study, light-activated Ga^3+^CHP was shown to be an efficient singlet oxygen producer (29). The singlet oxygen generated during aPDI pretreatment might affect the structure of bacterial surface proteins responsible for their attachment to the host (43). Moreover, a previous study showed that aPDI reduces biofilm adherence to abiotic surfaces (44). Our results showed for the first time that aPDI using green light-activated Ga^3+^CHP reduced the adhesion of *S. aureus* to a biotic surface, namely, keratinocytes (**Fig 6**). Interestingly, pretreatment with aPDI did not significantly change the level of internalization of *S. aureus*, regardless of inoculum size (MOI 10 vs. MOI 1) and aPDI strength (High vs. Low), and the number of intracellular *S. aureus* did not change (**Fig 6**). These results indicate that internalization is prioritized by *S. aureus* during infection until a certain capacity of host internalization is achieved. Finally, the bacterial inoculum also showed a lower proliferation rate after aPDI treatment, which we observed as a significant reduction in the growth of the extracellular fraction.

Some challenges need to be overcome for aPDI to be an anti-intracellular therapy (45). The ideal scenario is killing infected keratinocytes while sparing noninfected keratinocytes. To achieve this selective killing, the PS must accumulate inside the host cells and perhaps inside the intracellular vesicles where the pathogen may reside, which is a major challenge. For this reason, investigating the intracellular localization of both the pathogen and the PS is key to verifying the potential of aPDI treatment for intracellular pathogens. The second important issue is the effective cellular concentration of the PS, which depends not only on the accumulation process but also on the potential efflux of PS or the unwanted interaction of PS with host cell biomolecules. We studied Ga^3+^MPs and revealed divergent cellular localization patterns in keratinocytes. In general, the accumulation of porphyrins in eukaryotic cells occurs through a slow passive diffusion process with partial accumulation within mitochondria (45,46). In our study, cationic Ga^3+^CHP was mostly localized in lysosomal structures (**Fig 9**) and partially in mitochondria (**Fig S3B**). The presence of cationic quaternary ammonium moieties in the structure of Ga^3+^CHP might increase the targeting and uptake by lysosomes through electrostatic attraction (47,48). At this stage of research, it is difficult to determine whether Ga^3+^CHP first accumulates in lysosomes, attracting bacteria to these structures, or whether bacteria capture Ga^3+^CHP via highly specialized heme import systems, consequently ending up in lysosomes. On the one hand, the environment of the lysosome is nutrient-poor, which favors the bacterial cell in changing its phenotype to a dormant one and thus becoming more resistant to antimicrobials. On the other hand, in a poor environment, a pathogen such as *S. aureus* produces highly specialized proteins that efficiently capture nutrients such as heme or its structural analogs, as demonstrated in this work and our previous studies (**Fig 8**) (26,29). Interestingly, we did not confirm the lysosomal localization of intracellular *S. aureus* itself (**Fig 9**; *S. aureus* + lysosome). However, simultaneous colocalization of intracellular bacteria with Ga^3+^CHP in the lysosomal structures of the host was confirmed (**Fig 9**; *S. aureus* + Ga^3+^CHP + Lysosome). The presence of both *S. aureus* and Ga^3+^CHP may influence metabolic changes that favor the colocalization of *S. aureus* inside lysosomes. Therefore, there is a high probability that Ga^3+^CHP can reach intracellular bacteria or even be intracellularly accumulated through heme receptor acquisition systems as a part of the light-independent action of Ga^3+^MPs.

According to Strategy 3 presented in this study, we tested the effectiveness of aPDI on the intracellular load of *S. aureus* in keratinocytes. To date, there are a few studies on the effectiveness of aPDI in eliminating intracellular bacteria. For instance, efficient aPDI killing of intracellular *S. aureus* in HeLa cells was achieved with red light and a conjugate of the cell-binding domain of endolysin (CBD3) and silicon phthalocyanine (700DX) (49). Furthermore, blue light-activated gallium-substituted hemoglobin loaded on silver nanoparticles was used against intracellular *S. aureus*, which persisted inside professional phagocytes (50). Both examples use high-molecular-weight bioconjugates, which may have more difficulty accumulating inside eukaryotic cells than low-molecular-weight compounds. Previous studies on aPDI efficacy in this field have mainly focused on infection models involving either cancer cell lines or professional phagocytes, such as macrophages (49,50). Our study is the first evaluation of aPDI efficacy in eliminating *S. aureus* that reside in keratinocytes. The accumulation of bioconjugates may be difficult for nonprofessional phagocytes due to the lower accumulation capacity of these cells (51). We used small molecular weight compounds, gallium metalloporphyrins, rather than bioconjugation with large molecules. The addition of a positive charge gained by quaternary ammonium moieties increases the hydrophilicity of the compound, which might be important for PS lysosomal localization inside eukaryotic cells (52). Similar results were observed in the intracellular localization of hydrophilic sulfonated tetraphenyl porphyrins (53). Moreover, it has been reported that adding a cationic charge was necessary for effective photokilling and higher accumulation of benzophenoxazine in the intracellular pathogen *Leishmania* (54). Ga^3+^CHP was previously reported by us as a biocompatible and safe agent both *in vitro* and *in vivo* (30). Green light excitation of Ga^3+^CHP did not exhibit pronounced phototoxicity in studies on HaCaT cell cultures alone (29) or in our infectious model (**Fig 10**), as discussed below.

Previous studies on aPDI treatment of intracellular bacteria have mainly focused on intracellular photokilling efficacy without testing eukaryotic safety as a part of the infection model (49,50). Monitoring aPDI phototoxicity was focused on uninfected cell cultures separately rather than on infected cells harboring intracellular *S. aureus* (49,50). This approach does not reflect the actual impact of aPDI on infected cells, especially since the infection itself substantially reduces host cell growth. In our approach, we monitored growth in real time during the infection. At first Ga^3+^CHP was added and was incubated in the dark. This was followed by green light irradiation, after which the cells were cultured until they reached the plateau phase. After aPDI with Ga^3+^CHP, both infected and uninfected cells exhibited some phototoxicity reflected as a growth delay (**Fig 10 D vs. E**). Surprisingly, the growth rate of uninfected cells was relatively slower than that of infected cells, despite higher survival immediately after aPDI. Nevertheless, both infected and uninfected cells eventually reached a plateau phase, indicating that the photodamage could be overcome by the cell line. aPDI combined with Ga^3+^CHP is a promising anti-intracellular treatment that exhibits a low phototoxicity impact on keratinocytes despite the infection status. One limitation of this study is the inability to distinguish whether aPDI causes elimination of only intracellular bacteria without damaging keratinocytes or the bacteria together with the keratinocytes that incorporate them.

Overall, this research presents the application potential of light-activated compounds for i) the elimination of staphylococcal recurrent infections, ii) a reduction in infection severity and bacterial attachment to host cells, and iii) the elimination of intracellular *S. aureus*. We highlighted the great importance of the simultaneous colocalization of a PS and intracellular bacteria within the host cells to effectively reduce the number of infected cells by aPDI. This study revealed a variety of aPDI methods that can be used to overcome recurrent staphylococcal infections.

## Material and Methods

### Bacterial strains and growth conditions

In this study, we used two GFP-expressing *S. aureus* strains: hypervirulent USA300 (AH3369) derived from A. Horswill (55) and non-hypervirulent, *agr*-deficient RN4220 from BEI Resources, USA. Both strains were grown in trypticase soy broth (TSB, bioMérieux, France) at 37 °C with 10 μg/mL of either chloramphenicol for USA300 or trimethoprim for RN4220 to sustain the GFP plasmid. To obtain bacteria in the stationary phase of growth, overnight cultures were grown for 16-20 hours and diluted to 0.5 McFarland (∼10^7^ CFU/mL). For infection, stationary overnight cultures were diluted to 1:100 in flask and grown with shaking (150 rpm/37°C) to obtain logarithmic phase up to 2 hours then diluted to 0.5 McFarland for further investigations.

### Cell line and culture media (Antibiotic ON/OFF)

The human immortalized keratinocyte HaCaT cell line was used in this study. As the cell line for the filaggrin deficient cell line (*FLG* sh), we used knockdown cells after lentiviral particles FLG shRNA (sc-43364-V) infection, characterized previously (56). Control cells were treated with an empty vector (*FLG* ctrl). Culture medium with antibiotic (Antibiotic ON) consisted of Dulbecco’s modified Eagle’s medium (DMEM) with 10% fetal bovine serum (FBS), 4.5 g/L glucose, 1 mM sodium pyruvate, 100 U/mL penicillin, 100 μg/mL streptomycin, 2 mM l-glutamine, and 1 mM nonessential amino acids. The medium without antibiotic (Antibiotic OFF) was based on the previously given composition of DMEM HaCaT growth medium but without the addition of streptomycin and penicillin. All media components were derived from Gibco, Thermo Fisher Scientific, USA. Cells were grown in a standard humidified incubator at 37 °C in a 5% CO_2_ atmosphere.

**Figure 11.**
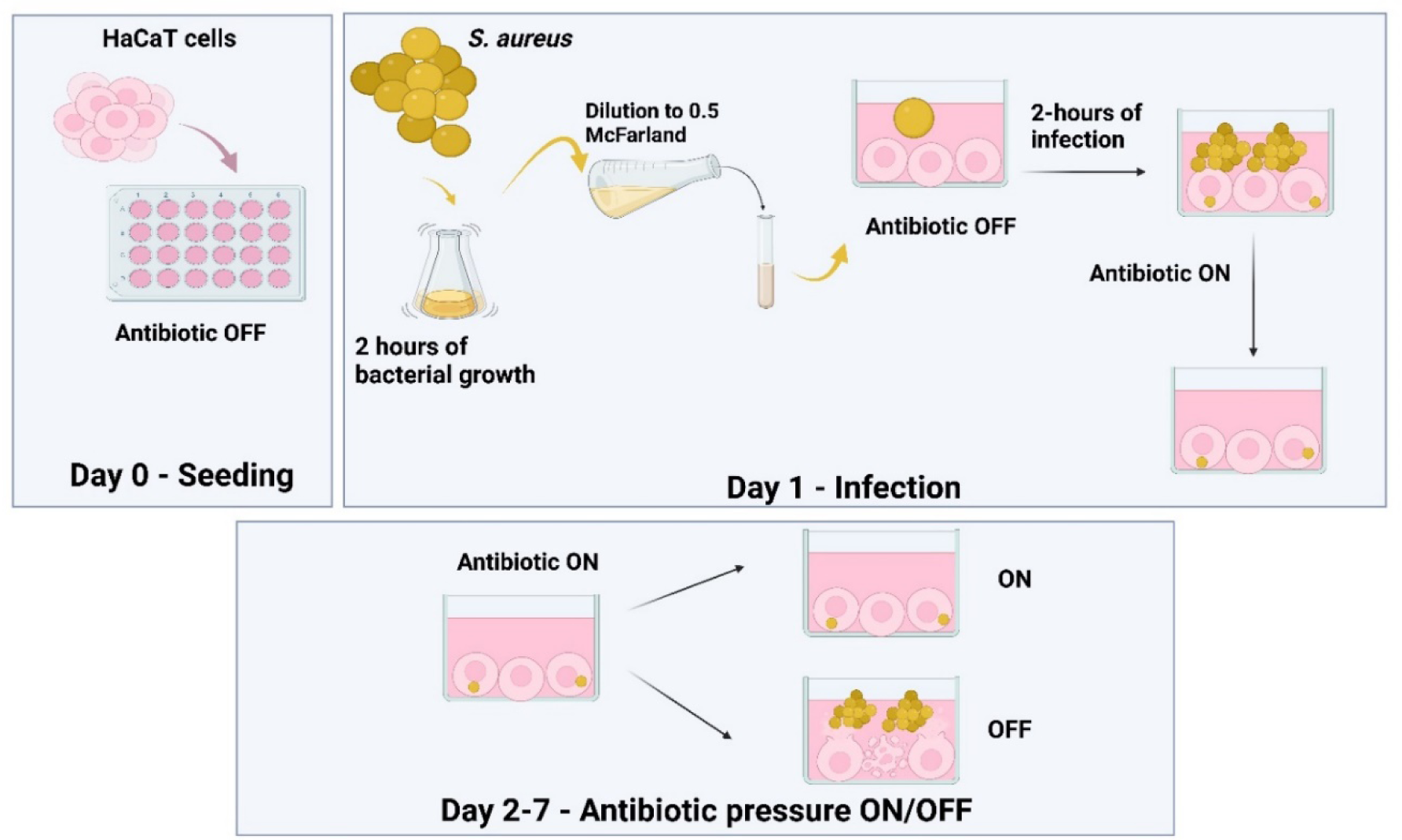
Methodology of the infection model preparation. On the Day 0, HaCaT cells were seeded at the 24-well plate in a non-antibiotic medium (Antibiotic OFF). On day 1, *S. aureus* overnight cultures were diluted and grown by 2 hours to achieve a logarithmic growth phase. Afterward, cultures were diluted and transferred to the cells at a proper multiplicity of infection. Then, after 2 hours of infection, the growth medium was changed to antibiotic (Antibiotic ON) to put antibiotic pressure on intracellular invasion. On days 2-7, the antibiotic pressure was maintained or lifted by changing the medium to non-antibiotic (Antibiotic OFF) to release extracellularly intracellular *S. aureus*.

### Co-culture methodology

To obtain intracellular *S. aureus*, we proposed the infection model protocol, provided in detail on **Fig 11**. Briefly, HaCaT cells were seeded in the antibiotic-free medium (Antibiotic OFF) prior to infection day, referred to as Day 0, at a density of 10^5^ cells/per well in a 24-well plate (Corning, USA). The next day (Day 1), overnight *S. aureus* cultures were diluted 1:100 in the 10 mL of TSB with proper antibiotic to sustain the plasmid in the flask. Cells were grown for 2 hours with 150 rpm shaking at 37°C to achieve logarithmic growth. Bacterial suspensions were diluted in TSB to 0.5 McFarland (∼2 x 10^7^ CFU/mL), and depending on the MOI for infection, the appropriate concentration of bacteria was added to the growth medium of HaCaT cells. After 2 hours of infection incubator at 37 °C in a 5% CO_2_ atmosphere, cells were washed with PBS, and the medium was changed to contain the standard cultures antibiotics (Antibiotic ON). During the following days (Day 2-7), either the antibiotic pressure was maintained, or the medium was changed for non-antibiotic (OFF) to release bacteria from the host.

To investigate the *S. aureus* fraction to count CFU/mL, we collected each fraction at the proper time point of the infection model preparation. For extracellular *S. aureus*, the fraction was collected from the growth medium after 2 hours of infection. For the intra+adherent fraction, after infection, cells were washed three times with PBS, and trypsinized with TrypLE^TM^ Express (Thermofisher Scientific, USA). Cells were collected, washed, centrifuged, then resuspended in the 0.1% TritionX-100 in MiliQ for 30 minutes for cell lysis. For intracellular *S. aureus*, 2 hours after the addition of *S. aureus* infectious inoculum, cells were washed, and the growth medium was change to fresh medium with antibiotic (Antibiotic ON) for kill the adherent and extracellular bacteria. After 1 hour incubation with antibiotics, the cells were washed and harvested. After that, samples were washed and lysed with lysis buffer. Each fraction was serially diluted and placed on TSA agar plate for CFU/mL counting.

To prepare the fraction of *S. aureus* from different stages of infection for analysis on the flow cytometer. After infection, cells were washed with PBS, then trypsinized and harvested. Culture medium was used to neutralize the trypsin, and the cells were washed with PBS and resuspended in BD Cytofix/CytoPrem^TM^ kit (BD Biosciences, USA). After a 20-minute incubation in the dark with pre-fix buffer, the cells were centrifuged, washed twice, and resuspended in PBS for further analysis on flow cytometer.

### Gallium compounds

The synthesis and structure characterization of cationic gallium porphyrin (Ga^3+^CHP) and Ga3^+^ mesoporphyrin IX chloride (Ga^3+^MPIX) was previously described in detail (29,30). The initial stock of Ga^3+^CHP was prepared in Milli-Q water, while Ga^3+^MPIX in the 0.1 M NaOH solution.

### Compound accumulation

The intracellular accumulation of each photosensitizer inside human keratinocytes was analyzed by two methods: flow cytometer analysis and measurements of fluorescence intensity of cell lysates. Ga^3+^MPIX or Ga^3+^CHP at 10 μM were added to the medium of HaCaT cells either infected or uninfected. After dark incubation at the desired time, the external photosensitizer was removed, then cells were trypsinized and collected. Cells were either fixed with BD Cytofix/CytoPrem^TM^ kit for flow cytometry analysis or counted and then lysed with 0.1M NaOH/1%SDS buffer for fluorescence lysate measurements. The fluorescence intensity of each sample was measured with an EnVision Multilabel Plate Reader (PerkinElmer, USA) at the following emission/excitation wavelengths: Ga^3+^MPIX at 406/573 nm and Ga^3+^CHP at 406/582 nm. Accumulation calculations for each PS were made from a compound calibration curve prepared in the lysis solution. The uptake values are presented as PS molecules accumulated per HaCaT cell number in the well. The molecular weight of Ga^3+^CHP was calculated to be 907.08 g/mol, and that of Ga^3+^MPIX was estimated to be 669.85 g/mol.

### Flow cytometry

Prepared fixed cells were analyzed by GFP signal originating from *S. aureus* that was detected with the green detector channel (exc 488 nm, ems 525/30 nm). For the accumulation of photosensitizer - the PS signal detected the Red Detector channel (exc 642 nm, ems 664/20 nm). Cells were analyzed using Guava easyCyte^TM^ flow cytometer and guavaSoft 3.1.1. software with an analysis of 10,000 events.

### Light activation of Ga^3+^MPs (Strategy 1, 2 and 3)

For the light-activation of gallium compounds, we used a light-emitting diode (LED) light emitting green 522 nm light (irradiance= 10.6 mW/cm^2^, FWHM= 34 nm) to excitement either Ga^3+^MPIX or Ga^3+^CHP. The excitement of gallium compounds within Q-bands at the emission spectra of lamp with the excitement of gallium compound was previously characterized in our previous research (57).

We applied light-activation of gallium metalloporphyrins on the infection model at different stages of the preparation, based on our proposed strategies (**Fig 2**). In strategy 1, HaCaT cells were infected with *S. aureus* at MOI 10 for 2 hours in an antibiotic-free medium (Antibiotic OFF). Then, cells were washed with PBS, and the medium was changed to antibiotic (Antibiotic ON). The next day, the medium was removed, adherent cells were washed with PBS, then the antibiotic-free medium was again applied (Antibiotic OFF) and cultivated for 16-20 hours to release the intracellular *S. aureus* from the host. Then, the medium was collected and centrifuged (10 000 rpm, 5 min), and bacterial sediment was resuspended in 1 mL of fresh TSB. The 10 µL of the bacterial sample was taken to count the number of bacteria released from the co-culture. The bacterial aliquot of 450 µL was mixed with 50 µL of 10x concentrated PS, then incubated in the dark with shaking at 37°C. The light illumination was applied, and surviving bacteria were serially diluted and placed on the TSA agar plates for CFU/mL counting.

For bacteria pre-treatment with aPDI at Strategy 2, the overnight *S. aureus* was diluted to 0.5 McFarland (10^7^ CFU/mL), then 900 µL of bacterial aliquots were placed on the 24-well plates with 100 µL of 10x concentrated photosensitizer, followed by 10 min dark incubation with shaking at 37°C. Then, samples were illuminated. For a low dose of aPDI, 1 µM of Ga^3+^CHP was used with 2 J/cm^2^ dose of green light (1 log_10_ of reduction in bacterial survival, final bacterial concentration of ∼10^6^ CFU/mL), while for a high dose, we used 1 µM of Ga^3+^CHP and 5 J/cm^2^ (2 log_10_ of reduction, final bacterial concentration: ∼10^5^ CFU/mL). Samples were centrifuged (10 000 rpm/5 min), then resuspended in 100 µL of fresh TSB before addition to the 10^5^ HaCaT cells to start the infection. Untreated cells were added at the proper MOI (10 or 1) as a corresponding control.

In strategy 3, either Ga^3+^MPIX or Ga^3+^CHP at 10 µM concentration was added to infected cells at the 1^st^ day post-infection. Then samples were left for dark incubation in the CO_2_ incubator for 2 or 6 hours. The PS-containing medium was replaced with PS-free medium for green light illumination at dose of either 6.36 J/cm^2^ or 12.72 J/cm^2^. Afterwards, cells were prepared for flow cytometry analysis. The same protocol was conducted for real-time growth analysis of the cells on the xCELLigence device, however, after illumination, cells were left for measurements of the Cell Index parameter.

### Real-time growth analysis

For real-time analysis of cell growth dynamics, HaCaT cells were seeded the day before at a density of 10^4^ per well on an E-plate (ACEA Biosciences Inc., USA) on the xCELLigence real-time cell analysis (RTCA) device (ACEA Biosciences Inc., USA). The next day, cells were treated as appropriate to the experimental purpose; either infection with proper MOI was conducted, or photosensitizer was added to the dark incubation studies. The Cell Index (X-axis) monitored the change in the flow of electrons between electrodes on the E-plate. The flow of impendence depends on the cell type, the density, its shape, and the degree of adhesion to the well (58). The CI parameter was monitored every 10 minutes until the plateau phase of the cell growth was reached.

### Confocal imaging

Specimens were imaged using a confocal laser scanning microscope (Leica SP8X equipped with an incubation chamber for the live analysis) with a 63× oil immersion lens (Leica, Germany). Cell nucleus for intracellular persistence studies and 3D-dimension images was stained with HOEST 33342 dye (Thermofisher Scientific, USA). For lysosomal staining the LysoTracker™ Deep Red (Sigma-Aldrich, USA) was used; for mitochondria – Mito RED (Sigma-Aldrich, USA); for the Golgi apparatus – GOLGI tracker NBD c6 ceramide (Thermofisher Scientific, USA). Pixel intensities were quantified and evaluated using the Pearson’s correlation or the overlap coefficient to derive the colocalization rate (%). Quantitative analyses of colocalizations were performed using Leica Application Suite X, version 3.5.2.18963.

### Statistics

Statistical analysis was performed using GraphPad Prism 8 (GraphPad Software, Inc., CA, USA). Quantitative variables were characterized by the arithmetic mean and the standard deviation of the mean. Data were analyzed using one-way or two-way ANOVA with Dunnett’s multiple comparison test.

## Acknowledgments

This work was supported by UGrants–start no. 533-0C30-GS28-23 (KS) funded by the University of Gdańsk. Funds for Statutory Activities for the Laboratory of Photobiology and Molecular Diagnostics, IFB of UG and MUG (531-N119-D098-23). The eukaryotic cell lines with suppression of filaggrin expression were kindly provided by Dr. Danuta Gutowska-Owsiak (Experimental and Translational Immunology Group). We express our sincere thanks to Dr. Andrea Lipińska for her valuable assistance with the flow cytometric analyses. We thank Dr. Agnieszka Bernat-Wójtowska for a constructive discussion on the infection model used.

